# Transposable Elements are differentially activated in cell lineages during the developing murine submandibular gland

**DOI:** 10.1101/2023.04.01.535217

**Authors:** Braulio Valdebenito-Maturana

## Abstract

The murine submandibular gland (SMG) is a model organ to study development, because it follows a branching morphogenesis pattern that is similar to that of lung, kidney, and other systems. It has been speculated that through its study, insights into regeneration and cancer could be obtained. Previously, using bulk RNA-Seq data, we reported that Transposable Elements (TEs) become activated during the SMG development. However, an outstanding question was as to whether their activity influenced different cell populations. Here, taking advantage of a single cell RNA-Seq atlas of the developing SMG, I studied TE expression to find out whether their activity can be recapitulated across its development, and if so, how they influenced cell types and cell fate specification. In this work, I found a total of 339 TEs that are markers of different cell populations, and then, through the modeling of the SMG development using Trajectory Inference methods, I found 2 TEs that could be potentially influencing differentiation processes. In sum, this short report reveals that TEs may be involved in the normal development of the SMG, and it highlights the importance of considering them in scRNA-Seq studies.

## Introduction

The murine submandibular gland (SMG) is the largest salivary organ accounting ~80% of saliva secretion. The SMG is an organ that follows a process of branching morphogenesis across its developmental stages, similar to that of the lung, mammary glands, kidney, vascular system, amongst others. Thus, the SMG is model organ to study development, with potential applications in regeneration and cancer therapies (1). The murine SMG development occurs throughout the embryonic (E) and postnatal (P) stages, with key stages at: ~E12, end bud formation from cell progenitors; ~E14, branching morphogenesis; ~E16, cell differentiation; ~P1, differentiation of acinar cells required for saliva secretion; ~P30, SMG becomes fully functional, and at the adult stage, complete acinar maturation has occurred (2).

Single cell RNA-Sequencing (scRNA-Seq) is a technological advance that allows the study of gene expression at individual cells. Unlike bulk RNA-Seq, which captures an average portrait of gene expression, scRNA-Seq allows the discovery of heterogeneity across tissues and organs. In turn, this technology has been successfully applied to study cellular diversity in cancer (3–5), regeneration (6), and in branching organs, including the murine SMG (1,7,8). Particularly, Hauser et al. recently published the scRNA-Seq atlas of the murine SMG development, comprising the E12, E14, E16, P1, P30 and adult stages. In their work, they characterized the gene expression profile of the different cell groups at each timepoint, and moreover, they modeled the developing differentiation processes through Trajectory Inference (TI) methods. TI is the estimation of cell decision and differentiation events from the use of single cell data, grouping and ordering cells across pseudotime based on their expression profiles (9). Amongst the key advantages of TI methods, is that they are unbiased, and thus allow the discovery of discovery of new cells responsible for particular decisions across the developmental model obtained (9). Despite all the insights gained with scRNA-Seq and its computational applications, Transposable Elements are still not routinely considered to further aid in the understanding of cell processes and cell heterogeneity, as it was in the case of the work by Hauser et al. It is thought that the study of these elements in scRNA-Seq works an further aid understand cell biology (10). Indeed, preliminary works have indicated that these elements can indeed be associated with specific cell types: a subset of mouse embryonic cells known as “2C-like” is marked by expression of the MERVL transposable element (11,12).

Transposable Elements (TEs) are repetitive elements, comprising about ~50% of both the human and mouse genomes (13). TEs have the ability to transpose either via a “copy-and-paste” mechanism (i.e., through reverse transcription of their mRNAs), as in the case of the *Long Terminal Repeats* (LTRs), *Long Interspersed Nuclear Elements* (LINEs) and *Short Interspersed Nuclear Elements* (SINEs); or via a “cut-and-paste” mechanisms (i.e., excision of the TE from its original genomic location and insertion elsewhere) as in the case of the DNA TEs (14). In mouse, the distribution of TEs is 41.9% SINE, 27.0% LINE, 26.6% LTR and 4.5% DNA (15). Across evolutionary timescales, most TEs have accumulated mutations that impede their functional activity (10). However, there is extensive evidence now that TEs can still become transcriptionally active, and throughout such transcription, they can perturb (up- or down-regulate) expression of nearby and/or distantly located genes (14).

We have previously analyzed TE expression in the murine SMG using bulk RNA-Seq. Briefly, we found that more than 9000 TEs became differentially regulated across the E14.5, E16.5, E18.5, P1, P28 and P70 timepoints (15). Moreover, through statistical association analysis, we predicted genes that could be potentially modulated by TEs. An outstanding issue from our previous work, was how TEs are expressed at the single cell level, and how they might affect the specification of cell populations. It has been speculated that TEs could have expression in specific cell groups, and thus be used as markers of these cells (10). Taking this into account, in this work, the following questions were addressed: (1) Are TEs expressed across the SMG development?, (2) Do TEs become activated in specific cell groups, and if so, what is their impact, and (3) How TE expression impact cell specification / differentiation?

## Methods

The scRNA-Seq dataset was previously published by Hauser et al. (1) and is publicly available at the Gene Expression Omnibus (GEO) database, accession code GSE150327. Raw FASTQ files were obtained using the SRA Toolkit (16). The *Mus musculus* mm10 genome version, along with its corresponding gene annotation in GTF format and the RepeatMasker repetitive element annotation, were obtained from the UCSC database (17). Reads were aligned to the genome using STAR (18), with options --*soloType CB_UMI_Simple,* to turn on scRNA-Seq alignment strategy, --*soloCBwhitelist,* to provide the cell barcode whitelist file, --*outSAMmultNmax 1,* to return at most 1 alignment for multimapped reads, and --outSAMattributes NH HI nM AS CR UR CB UB GX GN sS sQ sM, to add cell and gene info as SAM attributes.

Transposable Element expression was measured using SoloTE (12). Briefly, SoloTE adopts the following strategy: first, reads not having the GN tag (i.e., not assigned to a gene) are selected, and the overlap between these reads and TEs is assessed using BEDtools (19). All reads overlapping TEs are processed in the next step. These reads are subsequently classified into one of the following categories based on the number of mapping locations: *locus-specific,* if the MAPQ is equal to 255 (indicating unique alignment), or *subfamily-specific,* if the MAPQ was less than 255 (indicating multiple alignment). We have previously taken advantage of the usage of this metric in *Spatial Transcriptomics* data, which follows a similar computational process during alignment and demultiplexing, to that of scRNA-Seq (20). Although not all TE locus are retrieved, it still shows an improvement to the default strategy of directly summarizing expression at the subfamily level (12). SoloTE generates as final output an expression matrix, that contains both gene and TE expression per cell.

Downstream analysis of the scRNA-Seq count matrices generated with SoloTE was performed with Seurat v4.0.6 (21). First, all expression matrices were filtered to only keep the cells used in the original work, to ensure that only cells that met the quality control thresholds applied by Hauser et al. were used. Then, the standard Seurat protocol was used, which is comprised of the sequential application of the *NormalizeData, FindVariableFeatures, ScaleData, RunPCA, FindNeighbors, and FindClusters* functions. The specific parameters used for each function were based on the source code for the analysis in the original work, available at https://github.com/chiblyaa/Salivarv-Gland-Development/. Moreover, specific care was taken to ensure that the same number of cell clusters were obtained. Once all cell clusters across all timepoints were manually curated, the Seurat *FindAllMarkers* function was used to find TEs that could be preferentially expressed in specific cell groups. Finally, TEs having an average log_2_(Fold Change) ≥ 1 and adjusted P-value ≤ 0.05 were selected and analyzed.

Locus-specific analysis of TEs was carried out using BEDtools and the aforementioned UCSC gene annotation in GTF format. TEs located in introns or overlapping with exons, were associated with their respective host gene, whereas intergenic TEs were associated to their closest gene in the genome. This analysis was repeated for the locus-specific TEs detected at each stage, and the lists of genes obtained were used in PANTHER to obtain the biological pathways in which they are involved.

## Results

### TE expression across the scRNA-Seq atlas of the murine SMG development

To measure TE expression across the available scRNA-Seq murine SMG datasets, we performed read alignment to the *mm10 Mus musculus* genome, and the subsequent cell demultiplexing process, using STAR (Methods). STAR adds specific SAM tags to the alignment files, indicating whether a read is associated to a gene or no. Taking advantage of this characteristic, SoloTE was used to assess TE expression considering all reads not associated to genes (Methods). Based on the number of mapping locations, SoloTE reports the locus of TEs if they have uniquely mapped reads, whereas for non-unique reads, TE expression is summarized at the subfamily level. Although not ideal, this strategy still allows the retrieval of locus-specific TE expression, which in turn, might help to understand their potential regulatory impact on genes (12).

Upon measuring TE expression with SoloTE, the resulting matrices were analyzed with Seurat. This analysis mainly consisted in expression normalization, dimensional reduction, cell cluster identification and cell cluster marker identification (see Methods). The resulting cell clusterings obtained were manually curated to ensure consistency with those from the original work. Once this was confirmed, TE expression was studied. First, the overall TE expression of each of the 4 main TE types (LTR, LINE, SINE and DNA) was depicted in the UMAP dimensional reduction plots **(Figure** 1). Across the murine SMG development, we observe that all TEs are expressed, with SINEs being the predominant type. This in turn suggests that there could be potential gene regulation events mediated by TEs, as SINEs are well known to impact expression of genes, by influencing mRNA translation or degradation (22,23). Interestingly, LINES and LTRs were expressed also at the P30 and Adult stages. Although TE expression was thought to only occur at early developmental stages as consequences of global epigenetic derepression, and in disease and stress conditions, as consequence of cellular malfunction, we and others have shown that they indeed become transcriptionally active in both mouse and human healthy, adult tissues (20,24). Such reports suggest that TEs could have functional implications in normal and proper cell functioning. Thus, for the maturation and normal activities of the adult SMG they could be playing a similar role.

**Figure 1.**
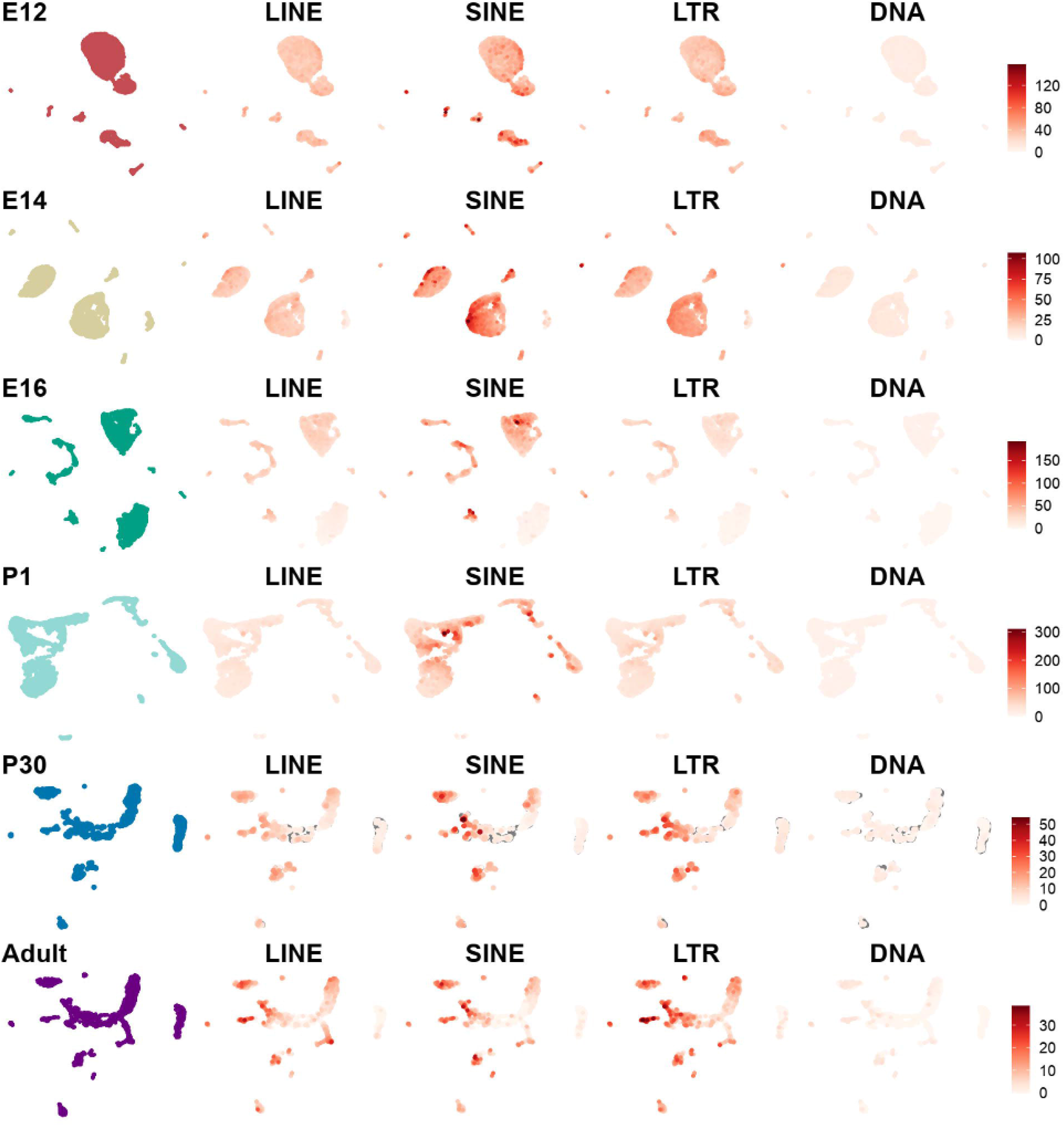
Overall TE expression across SMG development. First, the UMAP corresponding to each stage is presented and color-coded (E12: red, E14: dark yellow, E16: dark green, P1: light blue, P30: dark blue, Adult: purple), and then 4 UMAPs corresponding to the main TE classes (LINE, SINE, LTR and DNA) are depicted. The TE UMAPs show the number of UMIs of the respective TE class, according to the white-to-red scale indicated at the right.

As I was able to measure TE expression across the scRNA-Seq atlas of the murine SMG development, I then focused on the question of whether TEs are expressed preferentially in different cell groups, and thus, represent markers of specific cell types.

### TEs as cell markers in the murine SMG development

Upon the previous result, I then identified, annotated and manually curated the cell clusters per each timepoint. This was done to ensure that the same groups as in the original work were present in our analysis. Next, by using the Seurat function “FindAllMarkers”, the tables of genes and TEs significantly expressed in each group were generated. In turn, I was able to find between 27 – 79 TEs marking cellular clusters (Supplementary Table 1). Top TEs are indicated in **Figure 2** (E12), **Figure 3** (E14), **Figure 4** (E16), **Figure 5** (P1), **Figure 6** (P30) and **Figure 7** (Adult), with an overview in **Figure 8.** The marker TEs could be divided in two types: (1) High expression across a high proportion of cells in one cluster, and low expression across a low proportion of cells in the remaining clusters; (2) High expression across a high proportion of cells in one cluster, and lower expression across a high proportion of cells. The results of E12, E14, P1 and the Adult stages belong to the first group for the most part, whereas those from E16 and the P30 stages, have more markers belonging to the second group. This result reveals that for some TEs, expression is indeed restricted to specific cell types, whereas for other TEs, its activity is widespread (i.e., expression occurs across several clusters), but there are differential changes in the magnitude of expression in each one of them.

**Figure 2.**
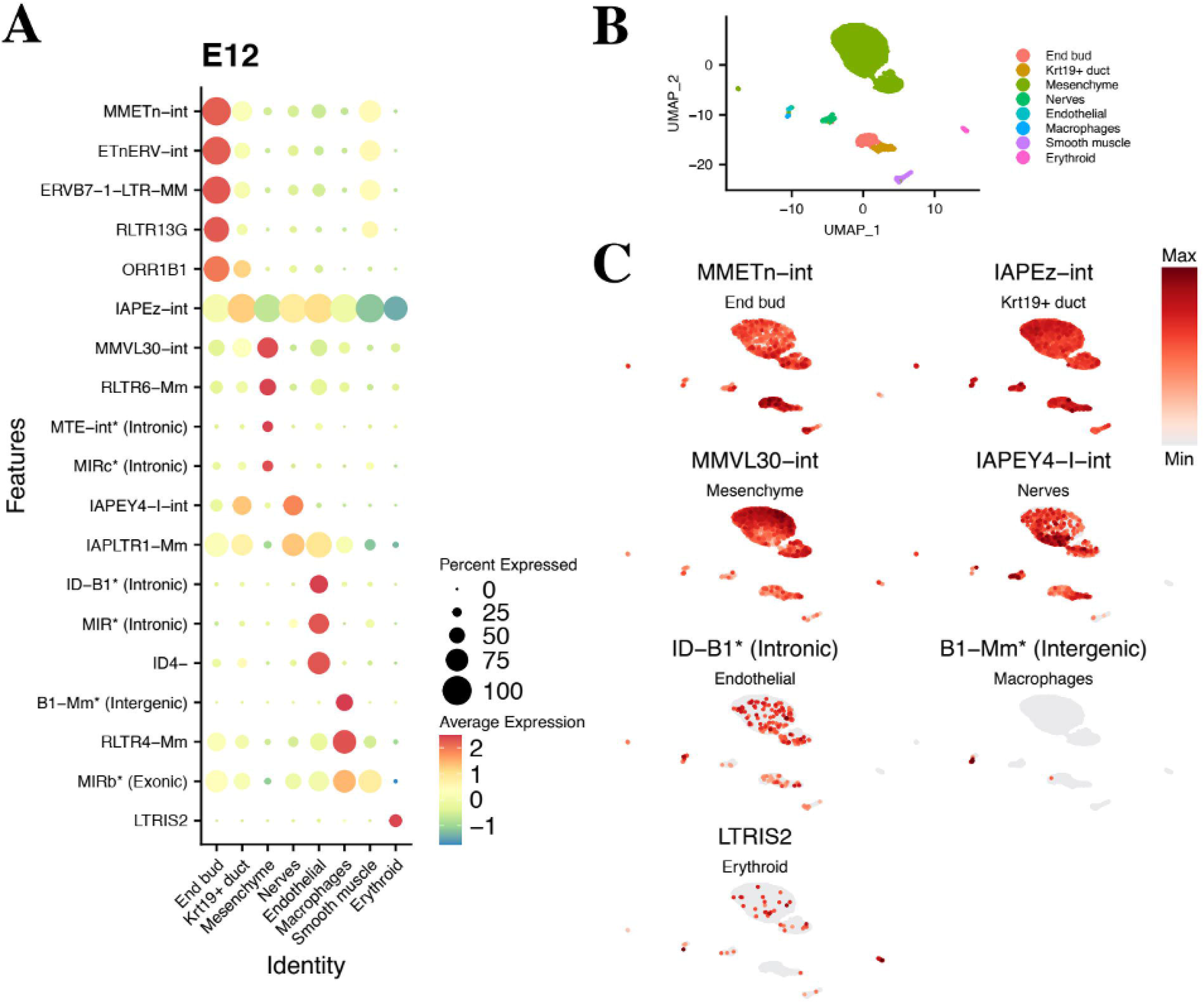
Selected marker TEs at E12. A. Dot plot of TEs (left) across cell clusters (bottom), indicating their average expression (color scale at the right) and percentage of cells per clusters in which they are expressed (size scale at the right). B. Reference UMAP indicating the distribution of each cell cluster. C. Expression in UMAP plot of the best marker TE of each cluster.

**Figure 3.**
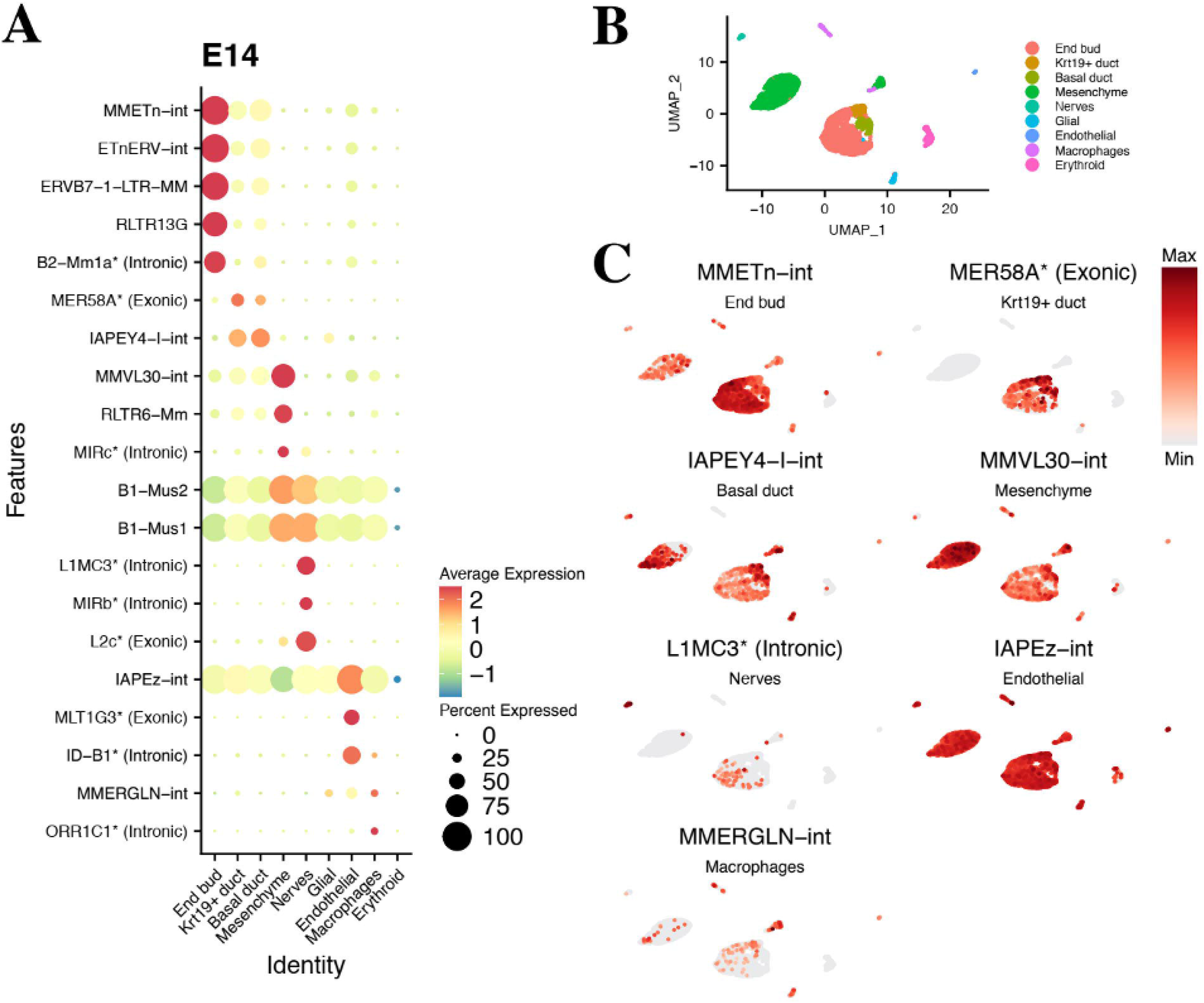
Selected marker TEs at E14. A. Dot plot of TEs (left) across cell clusters (bottom), indicating their average expression (color scale at the right) and percentage of cells per clusters in which they are expressed (size scale at the right). B. Reference UMAP indicating the distribution of each cell cluster. C. Expression in UMAP plot of the best marker TE of each cluster.

**Figure 4.**
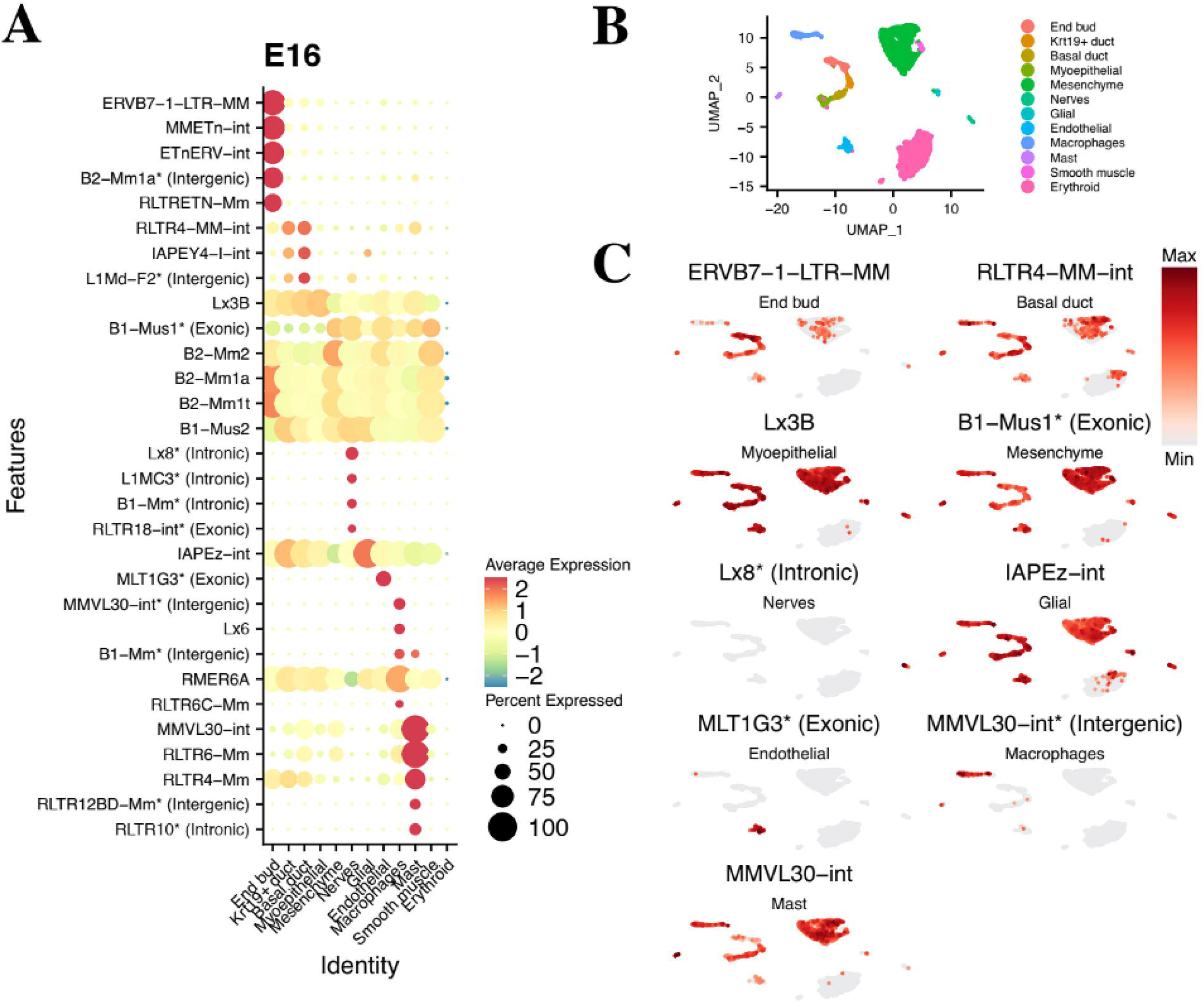
Selected marker TEs at E16. A. Dot plot of TEs (left) across cell clusters (bottom), indicating their average expression (color scale at the right) and percentage of cells per clusters in which they are expressed (size scale at the right). B. Reference UMAP indicating the distribution of each cell cluster. C. Expression in UMAP plot of the best marker TE of each cluster.

**Figure 5.**
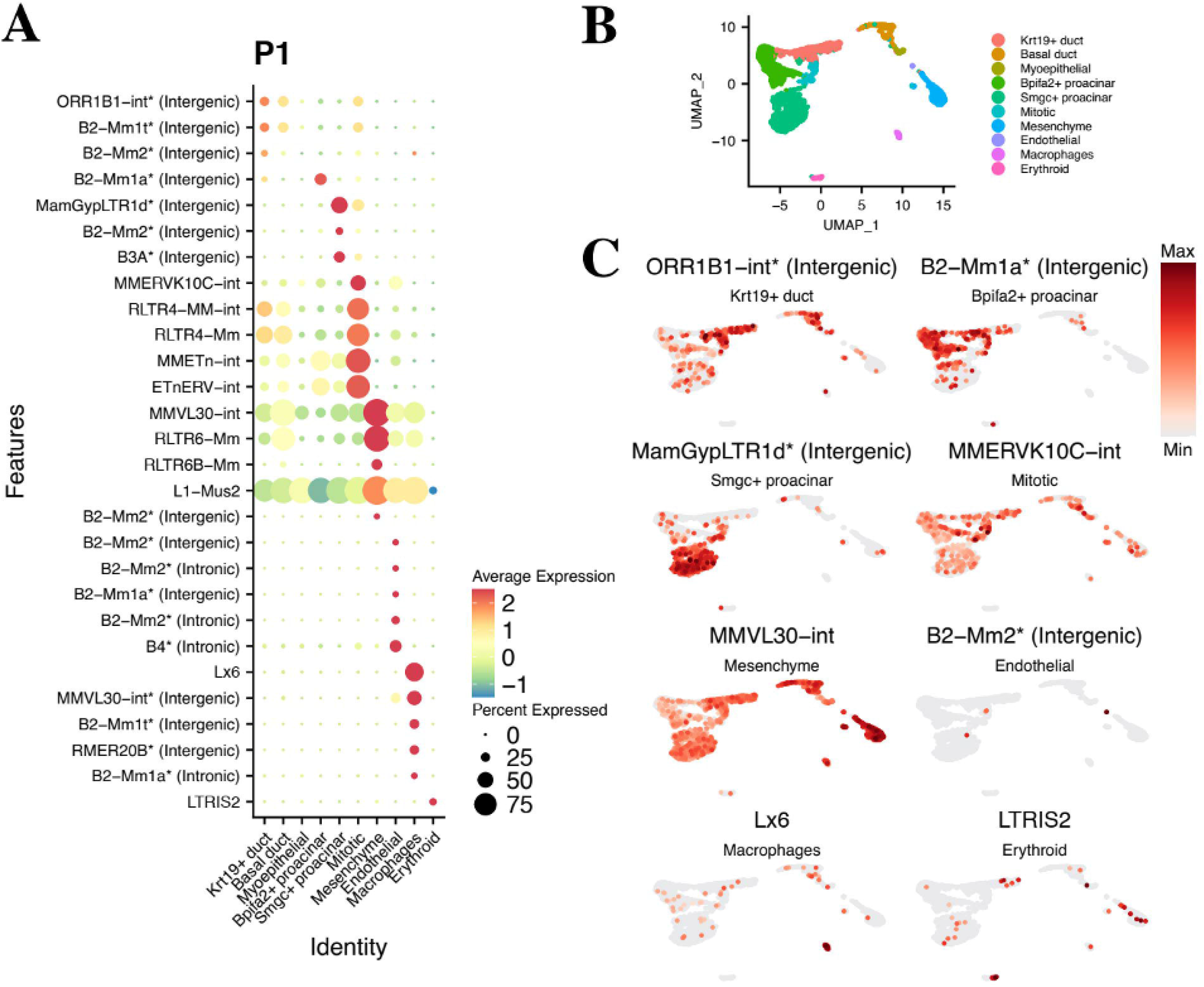
Selected marker TEs at P1. A. Dot plot of TEs (left) across cell clusters (bottom), indicating their average expression (color scale at the right) and percentage of cells per clusters in which they are expressed (size scale at the right). B. Reference UMAP indicating the distribution of each cell cluster. C. Expression in UMAP plot of the best marker TE of each cluster.

**Figure 6.**
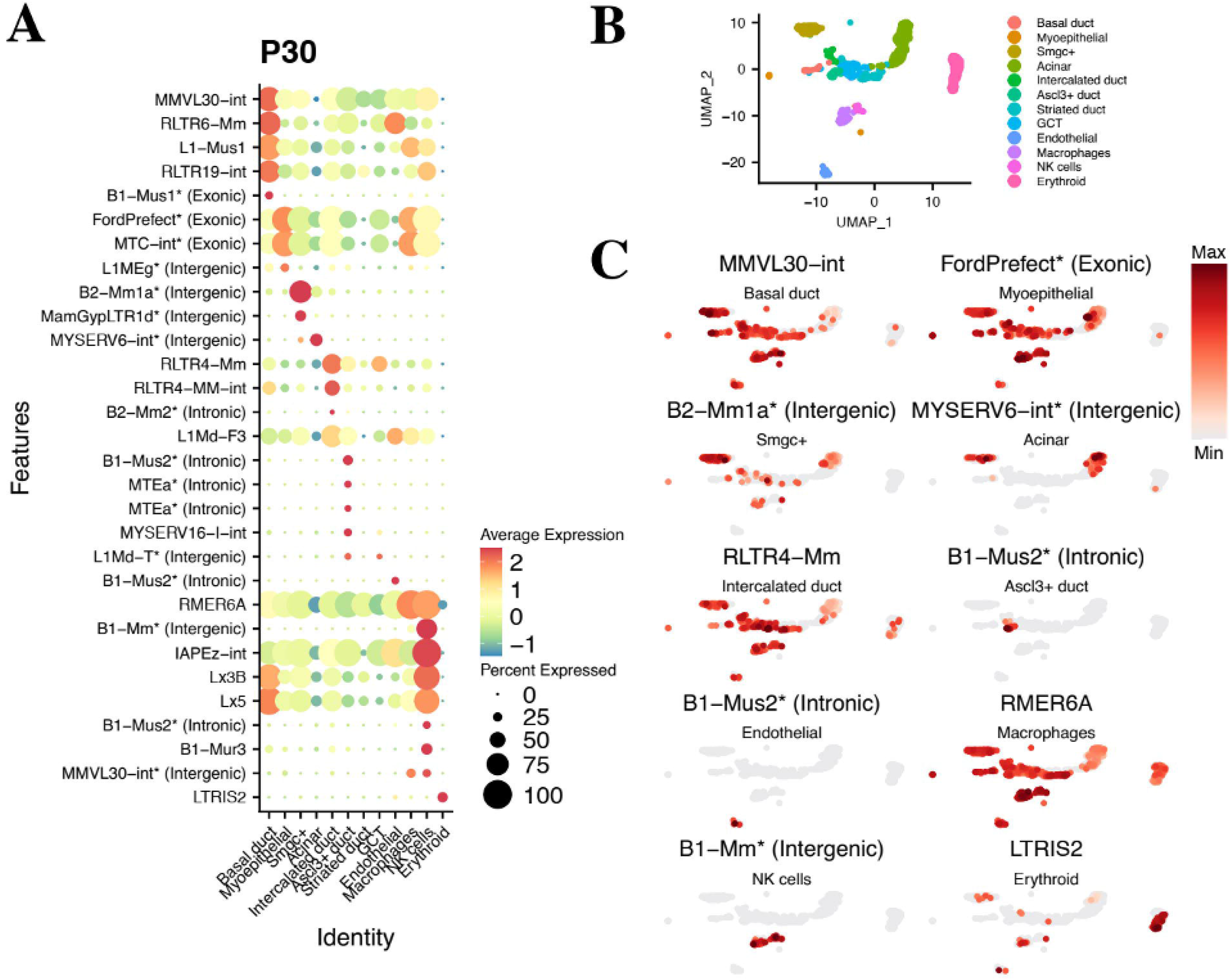
Selected marker TEs at P30. A. Dot plot of TEs (left) across cell clusters (bottom), indicating their average expression (color scale at the right) and percentage of cells per clusters in which they are expressed (size scale at the right). B. Reference UMAP indicating the distribution of each cell cluster. C. Expression in UMAP plot of the best marker TE of each cluster.

**Figure 7.**
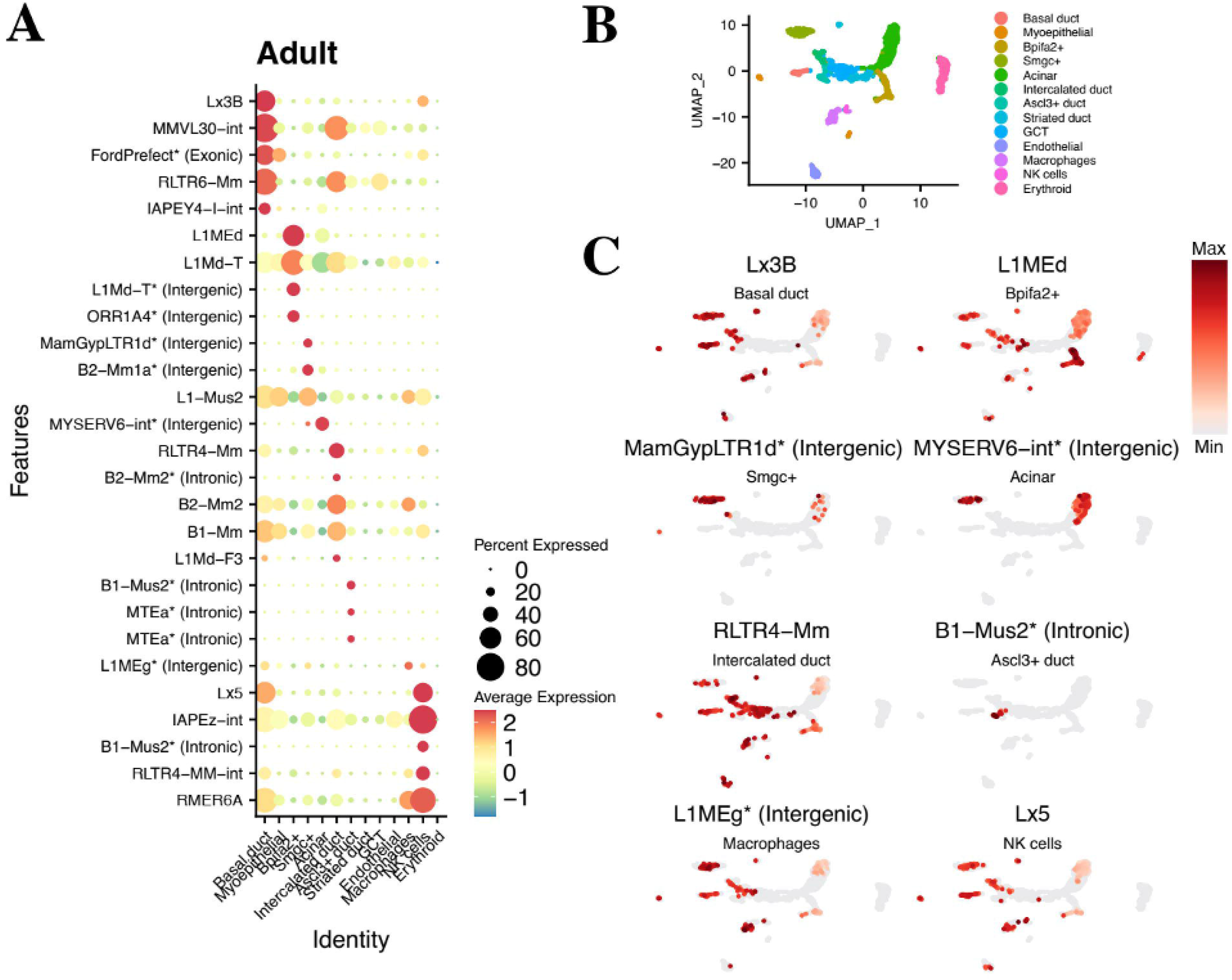
Selected marker TEs at Adult stage. A. Dot plot of TEs (left) across cell clusters (bottom), indicating their average expression (color scale at the right) and percentage of cells per clusters in which they are expressed (size scale at the right). **B.** Reference UMAP indicating the distribution of each cell cluster. C. Expression in UMAP plot of the best marker TE of each cluster.

**Figure 8.**
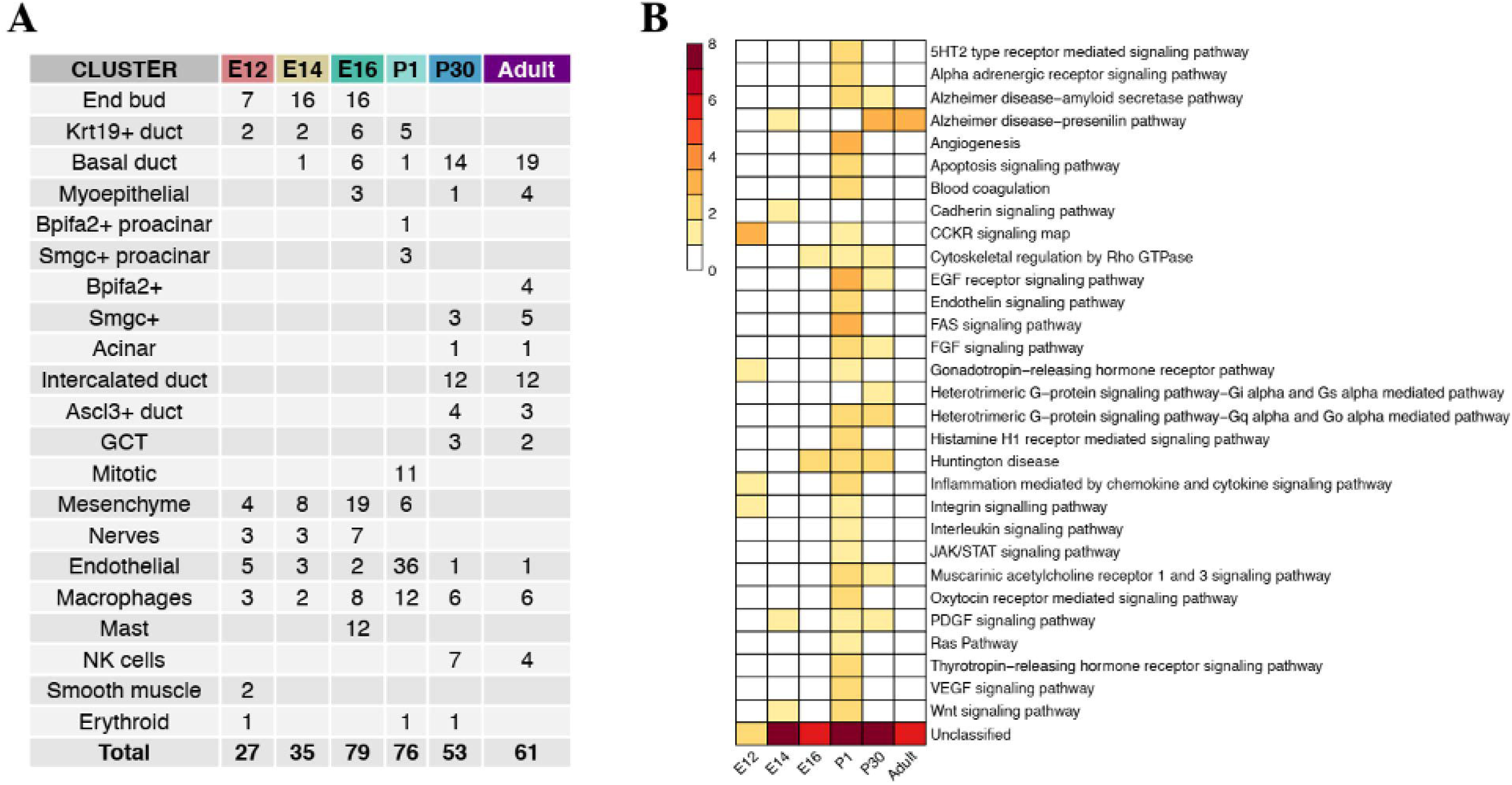
Overview of TE markers. A. Summary of marker TEs per each cell cluster (first column) across all timepoints (remaining columns). B. Heatmap of gene pathways associated to expressed TEs (color scale indicated at the left).

Based on the approach adopted by SoloTE to retain locus-specific information of TE expression, I also found that some marker TEs correspond to instances in particular genomic locations. Out of all TE markers detected, I was able to found between 11.4% - 75.0% of them corresponding to locus-specific TEs (Table 1). Although in all cases except P1, the proportion of these TEs is relatively low, I still highlight this result, due to its relevance to identify neighboring genes to those TEs, whose expression might be modulated by them.

**Table 1.** Locus-specific TE markers. The first column indicates the category (number of locus-specific TE markers or number of total markers), while columns 2-7 indicate the timepoint.

| **Category** | **E12** | **E14** | **E16** | **P1** | **P30** | **Adult** |
| --- | --- | --- | --- | --- | --- | --- |
| **Locus-specific TE markers** | 8 | 12 | 9 | 57 | 22 | 9 |
| **Total markers** | 27 | 35 | 79 | 76 | 53 | 61 |
| **Percentage of locus-specific markers** | 29.6% | 34.3% | 11.4% | 75.0% | 41.5% | 14.8% |

In turn, each of the locus-specific TE markers was associated to their closest gene, in order to predict their regulatory impact at the pathway level **(Figure 8),** following a similar approach to that of previous works (12,15,25,26). Most TE-related genes did not have an associated pathway, making it difficult to identify their potential consequences (Supplementary Table 2). On the other hand, amongst all pathways detected, I highlight the Integrin, Platelet-Derived Growth Factor (PDGF) and Wnt signaling pathways and Cytoskeletal regulation by Rho GTPase for E stages, and Apoptosis, Cytoskeletal regulation by Rho GTPase, along with the Epidermal Growth Factor (EGF), Fibroblast Growth Factor (FGF) and PDGF signaling pathways, for P stages. The Wnt signaling pathways has been implicated in salivary gland morphogenesis, whereas the Integrin signaling and the Cytoskeletal regulation pathways could be linked to the results previously reported by Gluck et al. (27) in their bulk RNA-Seq murine submandibular gland atlas, collectively suggesting a potential role of TE-mediated regulation. In addition, Cytoskeletal regulation by Rho

GTPase has been reported to play a key role in migration of cells in the salivary gland of the fruit fly *Drosophila melanogaster* (28). Thus, the putative interaction of TEs with genes acting in that pathway could also be of significance to the murine SMG. Interestingly, current evidence pinpoints to a role of PDGF coupled with the FGF signaling pathway in SMG branching morphogenesis and also in the interaction that occurs between mesenchymal and epithelial cells (29). Of note, the TE found linked to the PDGF pathway is indeed a marker of the mesenchymal cluster at E14, further supporting this notion. At P1 endothelial cells, TE markers linked to angiogenesis were found. Angiogenesis is a key process for the maintenance and localization of endothelial cells and evidence points to a role of these cells in modulating the patterning of the salivary gland during its development (30). Finally, the EGF signaling pathway has also been implicated during early SMG (31), and now it is well accepted as an important element of development, playing a wide spectrum of roles such as cell proliferation, migration, differentiation and apoptosis (32,33). Taking all of the above together, a novel and potential role of TEs in several key pathways for SMG could be suggested. Additionally, these results provide a preliminary answer to the original question of whether TEs can act as cell markers, along with a projection of their impact in these cell groups.

### Pseudotemporal TE expression in the murine SMG epithelial cells

The sorting and grouping of cells along a pseudotemporal developmental trajectory, as inferred with Trajectory Inference (TI) methods, can help understand cell specification, differentiation and branching events. To get insights on the role of TE expression in such cell processes, across SMG development, TI was following using the same guidelines used in the original work. Briefly, the PAGA Tree method implemented in the Dynverse R package (9) was used, and the E12 cells were defined as the starting point of the trajectory. Moreover, I only focused in epithelial cell populations as I noted that the usage of all cell types introduced artifacts, which was also reported in the original work. Once the TI method was applied, first I assessed the overall impact of the inclusion of TEs in the TI step. To this end, I compared the timepoint of cells in the trajectory to their pseudotime score **(Figure 9A** and **Figure 9B,** respectively). I observed that for the most part, cells with lower pseudotime scores correspond to cells of early stages, and cells with higher pseudotime scores to cells of the later stages. In addition, the cell type trajectory **(Figure 9C)** also is similar to the one shown in the original work. Collectively, these results indicate that the modelled trajectory is in agreement with the time-course progression of the SMG development.

**Figure 9.**
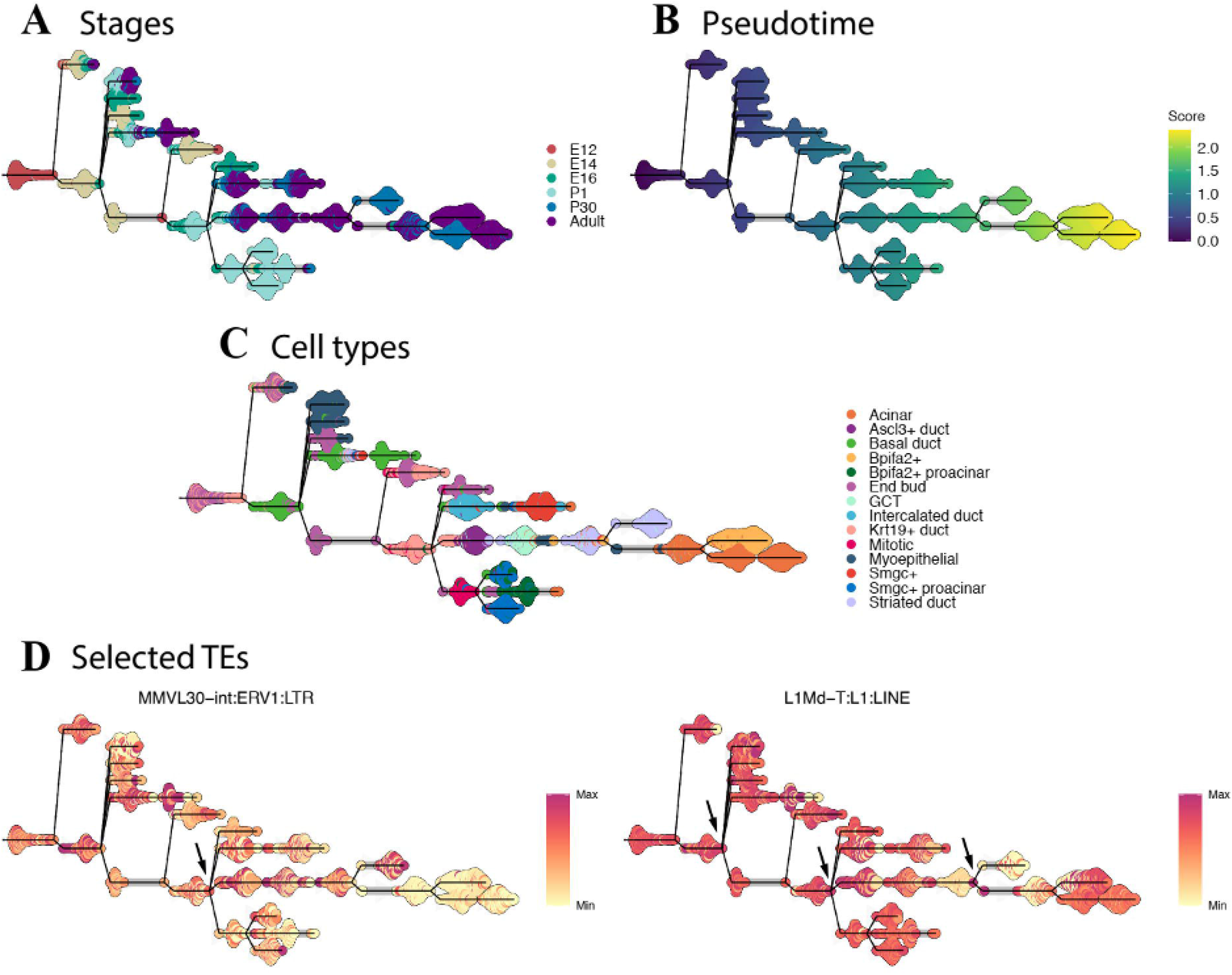
TE time course expression using Trajectory Inference (TI). TI colored by stage. **B.** TI colored by pseudotime score **C.** TI colored by cell types. **D.** TI of selected TEs, using a scale from light yellow to red, corresponding to its relative expression across the trajectories. Arrows indicate branching point cells in which TE expression is higher, and could potentially impact cell fate specification.

Afterwards, I focused and studied only a subset of TEs in the pseudotemporal developmental trajectory. From the previous cell marker analysis, I assessed the number of timepoints each TE appeared. Then, TEs expressed in either 5 or all of the 6 stages were selected, resulting in a total of 5 TEs. To understand the cell populations in which TEs could potentially be acting, I then plotted TI dendrograms colored by the expression of 2 TEs (MMVL30-int:ERV1:LTR and L1Md-T:L1:LINE) which were the most representative. Those TEs could be divided in two categories depending on their expression profile across the pseudotemporal trajectory: high expression on the cells appearing early in the dendrogram **(Figure 9D,** left), and high expression in cells appearing later in the dendrogram **(Figure 9D,** right). For the MMVL30 TE, there is higher expression in a branching Krt19+ duct cell **(Figure 9D,** left, arrow). This is a key branching point, because, and also noted originally by Hauser et al., all postnatal cell types appear from that point onwards, with the exception of Basal duct and Myoepithelial types that branch off early in the trajectory. Interestingly, the other TE, L1Md-T, could be acting in the differentiation towards the Basal duct and Myoepithelial cells, as it has higher expression in the branching point from which they derive **(Figure 9D,** right, first arrow). Moreover, the second point in which this TE has high expression is the same as the MMVL30 TE **(Figure 9D,** right, second arrow), thus further highlighting their putative influence in cell specification processes related to the postnatal populations. The last branching point in which this TE could be active **(Figure 9D,** right, third arrow) comes before the Striated Ducts and Acinar cells appear, which are the most prevalent cell groups in the adult murine SMG (34). Collectively, these findings can potentially suggest that TEs might be playing an important role in cell specification processes, both in the early developing SMG and during its last maturation steps.

## Discussion

We previously published the locus-specific analysis of TEs across the SMG development using bulk RNA-Seq data. One of the outstanding questions from that work, was the potential influence of TEs in SMG development at the single cell level. Here, I sought to address that question using a recently published scRNA-Seq atlas.

Unlike bulk RNA-Seq, there is only 1 tool published for the analysis of TEs in scRNA-Seq data, and it omits the genomic location of expressed TEs. This is a key point, because it its known that TEs can influence neighboring gene expression. In previous works, I and others have seen that by exploiting alignment metrics, such as the MAPQ, the loci of expressed TEs can still be retrieved. This is because in the mouse genomes most TEs correspond to evolutionary old TEs, and thus, have enough mutations that render them different from other copies. Thus, in this work, using SoloTE, which adopts a similar strategy, some of the locations of expressed TEs can still be retrieved, which might be useful in order to study their putative influence in genes located nearby. Indeed, from the cell marker analysis, I was able to found some TEs whose expression comes uniquely from one genomic location, further highlighting that particular instances might be playing important roles in cell groups.

Another potential limitation of this study is that 10X Genomics scRNA-Seq experiments capture poly-adenylated (polyA) transcripts. This was also a concern in our previous bulk RNA-Seq study, where the RNA selection protocol was based on polyA selection. In turn, we demonstrated that under these conditions, approximately ~70% of all TEs could be retrieved (15). Interestingly, another reports have revealed a similar finding, in that such polyA-selection protocol might hinder detection of other repetitive elements, such as satellites, but not of TEs (35,36). Thus, as I still obtained important and novel findings with using polyA transcripts, I speculate that the role of TEs could probably be higher if the full spectrum could be captured in sequencing experiments.

## Conclusions

scRNA-Seq, as the bulk RNA-Seq predecessor technology, is revolutionizing the impact and extent of gene expression studies. Although many technical advances have been made to study TEs in bulk RNA-Seq, they are not routinely studied, and scRNA-Seq suffers from this same pitfall. As TEs could play key role in modulating gene expression, it is thought that they can influence cell phenotypes and cell heterogeneity. Thus, their incorporation to scRNA-Seq studies can help further elucidate their role in important cellular processes such as cell differentiation.

Here I took advantage of a previously published scRNA-Seq atlas of the murine SMG development, and analyzed TE expression. I found that TEs are widespread expressed, and that some TEs have expression restricted to specific cell types. Moreover, I found TEs that could potentially be impacting cell decision events, further highlighting the importance of considering them in scRNA-Seq studies.

